# The SYP123-VAMP727 SNARE complex is involved in the delivery of inner cell wall components to the root hair shank in Arabidopsis

**DOI:** 10.1101/2020.12.28.424500

**Authors:** Tomoko Hirano, Kazuo Ebine, Takashi Ueda, Takumi Higaki, Takahiro Nakayama, Hiroki Konno, Hisako Takigawa-Imamura, Masa H. Sato

## Abstract

A root hair is a long tubular protrusion from a root hair cell established via tip growth, which is accomplished by the polarized deposition of membranous and cell wall components at the root hair apex accompanied by simultaneous hardening of the shank. The polarized secretion of materials to the root hair apex is well investigated; however, little is known about the deposition of inner cell wall materials at the root hair shank. We have previously reported that phosphatidylinositol-3,5-bisphosphate (PtdIns(3,5)P_2_)/ROP10 signaling is required for the regulation of cortical microtubule construction and the deposition of inner cell wall components at the root hair shank during hardening. To unravel the alternate secretion mechanism for delivery of the inner cell wall components to root hair shank, here, we demonstrate that root hair-specific Qa-SNARE, SYP123, localizes to the subapical zone and shank of elongating root hairs in Arabidopsis. SYP123-mediated root hair elongation was inhibited by the FAB1 inhibitor YM201636, and inhibition of PtdIns(3,5)P_2_ production impaired the plasma membrane localization of SYP123. We also showed that SYP123 forms a SNARE complex with VAMP727 on the plasma membrane, and *syp123* and *vamp727* mutants exhibited lower cell wall stiffness in the root hair shank because of impaired deposition of inner cell wall components. These results indicate that SYP123/VAMP727-mediated secretion is involved in the transport of inner cell wall components for hardening of the root hair shank.

## Introduction

The trafficking of proteins to various cell compartments by vesicular transport is essential for the viability of all eukaryotic cells. Transport vesicles carry cargo proteins from one compartment to another and discharge their cargo into specific compartments through membrane fusion. The specificity of membrane fusion is mainly mediated by RAB-GTPases ^1^ and soluble N-ethylmaleimide-sensitive factor attachment protein receptors (SNAREs), which comprise a large superfamily in eukaryotes ^2^. SNAREs have been categorized into target and vesicle SNAREs based on their functional localization, and are designated Q (glutamine)- or R (arginine)-SNAREs based on the conserved amino acid residues within their SNARE motif ^3,4^. Q-SNAREs are further subdivided into Qa (syntaxin)-, Qb-, Qc-, or Qbc-SNAREs based on their homology with synaptic SNAREs. When membrane fusion occurs between the target membrane and a transport vesicle, a specific set of target membrane SNAREs (Qa-, Qb-, Qc- or Qa-, Qbc-SNAREs) and vesicle SNAREs (R-SNAREs) form a SNARE complex at the membrane fusion site ^5^. As a specific set of the SNARE complex is involved in a particular membrane fusion event in each membrane trafficking pathway, specific cell compartments are marked by a specific set of SNARE complexes in their membrane ^6^.

During evolution, higher plant cells have developed various unique features, mostly in intracellular compartments and membrane trafficking systems ^7^. Membrane fusion events have been proven to function in various processes, including tissue development, signal transduction, physiological responses, and polarized secretion at the plasma membrane (PM) ^8^. Thus, higher plants have a more complex membrane trafficking system than other higher eukaryotes to establish the complex trafficking events, and land plants have more SNARE-encoding genes than unicellular plants ^9^. The increase in endosomal SNARE genes might be associated with plant cell development. In particular, a large number of Qa-SNAREs, which belong to the SYP1 family of SNAREs, are localized on the PM in Arabidopsis ^10^. SYP1s are further divided into three subfamilies: SYP11 (SYP111/KNOLLE, SYP112), SYP12 (SYP121/PEN1, SYP122, SYP123, SYP124, SYP125), and SYP13 (SYP131, SYP132). These SYP1 syntaxins show different spatiotemporal expression ^11^ and have different physiological functions. For instance, SYP132 is expressed ubiquitously in all tissues and uniformly localizes to the PM, suggesting that it functions in non-polar *de novo* secretion of PM materials in various tissues in Arabidopsis ^11,12^. SYP13-containing SNARE complexes are found in the charophycean alga *Klebsormidium flaccidum;* thus, SYP132 is thought to be evolutionarily ancient, and it is highly conserved in land plants ^13^. A new branch of SYP12 family SNAREs occurred in early land plants, such as the moss *Marchantia polymorpha* ^14^. The SYP12 family of genes expanded by gene duplication, and duplicated genes acquired specialized functions in various steps of secretion events during land plant evolution ^9,13,14^.

Arabidopsis VAMP/R-SNARE family proteins, VAMP72s (VAMP721/722/724/725), are generally localized to the trans-Golgi network and the PM and function in the secretory pathway ^10,15^. However, VAMP727 localizes to the plant-unique RAB5, ARA6, or FAB1-positive late endosomes ^16–18^. VAMP727 mediates not only membrane fusion between the pre-vacuolar compartment and the vacuolar membrane by forming a SNARE complex with SYP22, VTI11, and SYP51 ^16^, but also secretion via late endosomes by forming a complex with SYP121/PEN1 ^19^. This suggests that plants possess a unique late-endosomal trafficking pathway to the PM in addition to the conventional trans-Golgi network-PM secretion pathway mediated by VAMP721/722.

Root hairs are fast-growing tubular protrusions on root epidermal cells that play important roles in water and nutrient uptake in plants ^20^. The tip-focused polarized growth of root hairs is accomplished by the secretion of newly synthesized materials to the root hair tip via a polarized membrane trafficking mechanism. In Arabidopsis, two PM Qa-SNAREs in root hair cells, SYP123 and SYP132, function in root hair growth. SYP123 localizes to the tip region of root hairs by recycling between brefeldin A-sensitive endosomes and the PM of the expanding tip in an F-actin-dependent manner, whereas SYP132 is involved in non-polar *de novo* secretion. SYP123 and SYP132 act in a coordinated fashion along with transport vesicle-associated R-SNAREs VAMP721 and VAMP722 to mediate tip-focused membrane trafficking for root hair tip growth ^12^. Root hair formation is accomplished not only by tip growth, but also by simultaneous hardening of the shank through the production of an inner cell wall layer to form an elongated tubular structure ^21,22^. We have previously shown that phosphatidylinositol-3,5-bisphosphate [PtdIns(3,5)P_2_], the enzymatic product of FORMATION OF APLOID AND BINUCLEATE CELLS 1 (FAB1), is involved in root hair shank hardening in Arabidopsis. FAB1 and PtdIns(3,5)P_2_ localize to the plasma membrane along the shank of growing root hairs, whereas phosphatidylinositol 4-phosphate 5-kinase 3 (PIP5K3) and PtdIns(4,5)P2 localize to the root hair apex, where they are required for tip growth. A reduction in FAB1 function leads to wavy instead of straight root hairs, indicating that root hair shank hardening requires PtdIns(3,5)P_2_/ROP10 signaling. These results suggest the presence of an alternate secretion pathway for the delivery of inner cell wall components for shank hardening ^23^. In this study,we tried to identify the alternate secretion mechanism for delivery of the inner cell wall components and demonstrated that the root hair-specific Qa-SNARE protein, SYP123, in the root hair shank is involved in the transport of inner cell wall components for shank hardening.

## Results

### SYP123 localizes to the subapical zone and shank, but not the apex of elongating root hairs

We previously reported that SYP123 localizes to the apex region of root hairs by recycling between brefeldin A-sensitive endosomes and the PM of the expanding root hair tip ^12^. SYP124 and SYP125, the closest paralogs of SYP123, are localized to the PM of the shank or the subapical region, but not the apex, of the growing pollen tube ^24–26^. In the present study, we re-examined the dynamics of the PM localization of green fluorescent protein (GFP)-SYP123 in elongating root hair. We found that GFP-SYP123 localized to the PMs of the subapical region and the shank of the root hair, but not the root hair apex, whereas cytosolic GFP-SYP123 was observed in the apical region of the growing root hair (Fig. 1 and *SI Appendix*, Fig. S1, Movie S1). PM localization of GFP-SYP123 in the tip region was observed only after root hair elongation was terminated (*SI Appendix*, Fig. S1, Movie S2). The localization pattern of GFP-SYP123 during root hair elongation was similar to that of FAB1 and its product PI(3,5)P_2_ in root hair cells ^23^, suggesting the involvement of FAB1 and/or PtdIns(3,5)P_2_ function in the root hair shank localization of SYP123.

**Fig. 1.**
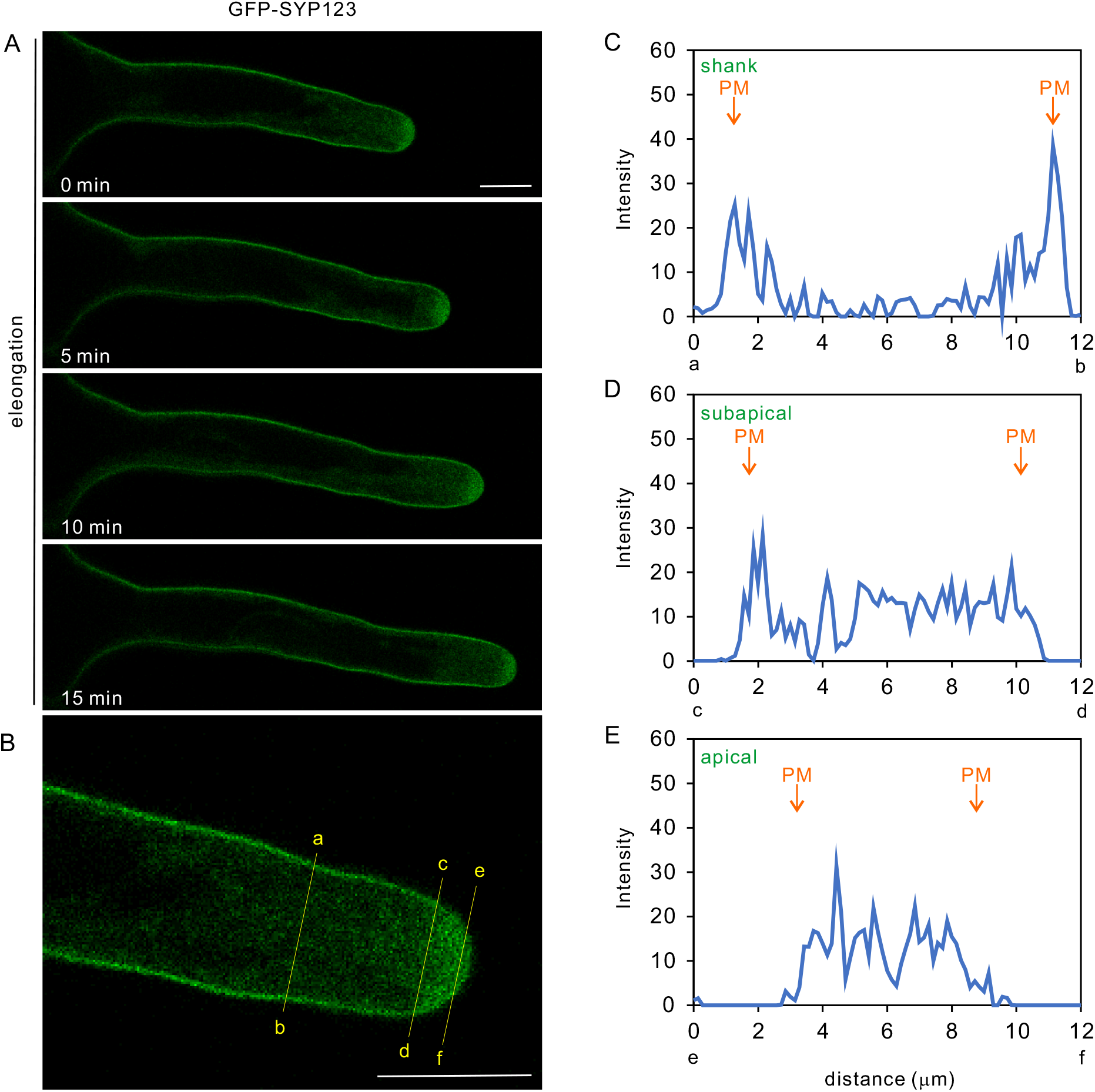
GFP-SYP123 predominantly localizes to the PM in the root hair shank in the elongating root hair. (A) Time-lapse images of an elongating root hair. The elongating root hair was observed by confocal laser microscopy in 5-min intervals. (B) Magnified images of the apical region of the elongating root hair. Fluorescence intensity profiles of GFP-SYP123 in the root hair shank in the [(a)–(b)] subapical [(c)–(d)] and apical [(e)–(f)] regions of an elongating root hair, as indicated in (B). Bars = 10 μm.

### PtdIns(3,5)P_2_ is involved in SYP123-dependent root hair elongation

In Arabidopsis, SYP123 and SYP132 are involved in distinct membrane trafficking pathways at the PM of root hairs: SYP123 is involved in the brefeldin A-sensitive recycling pathway in the apex region, whereas SYP132 acts in non-polar secretion throughout the root hair ^12^. We previously reported that root hair elongation is significantly inhibited by knockdown of *FAB1* or inhibition of PtdIns(3,5)P_2_ synthesis ^23,27^. In the present study, we investigated whether PtdIns(3,5)P_2_ is involved in SYP123-dependent root hair elongation by using a specific inhibitor for PtdIns(3,5)P_2_ synthesis, YM201636 ^17,28^.

In wild-type (WT) plants, the root hair length decreased by 40% in the presence of 1 μM YM201636. The root hair length of the *syp123* mutant was similar to that of WT plants in the presence of 1 μM YM201636, but was unchanged by incubation with YM201636, suggesting that SYP123-dependent root hair growth is sensitive to PtdIns (3,5)P_2_ inhibition (Fig. 2A). Further, root hairs of the *syp123* mutant were significantly broadened (Fig. 2B). The average surface area, length, and diameter of root hairs were 16.67 □m^2^, 526.1 ± 95.06 □m (n = 200) and 10.43 ± 2.45 □m (n = 30), respectively, for WT, and 16.38 □m^2^, 317.8 ± 97.3 □m (n = 200), and 17.31 ± 4.46 □m (n = 30), respectively, for *syp123*. This indicated that the surface area was unchanged by impairment of SYP123 function.

**Fig. 2.**
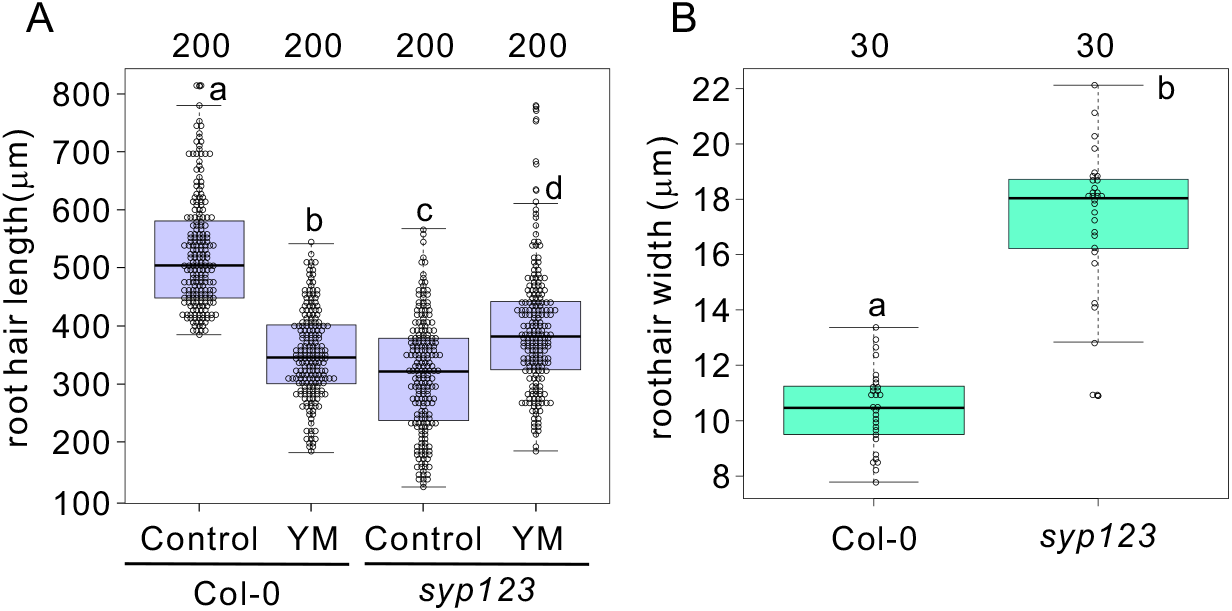
The *syp123* mutant has shorter and broader root hairs than the WT and is insensitive to YM201636. Measurement of root hair length (A) and width (B). Significant differences (*P* < 0.01; indicated by different letters) among genotypes were determined for each condition by one-way ANOVA followed by Tukey’s test (A) or the Mann-Whitney U-test (B). In the box- and-whisker plots, the boxes and solid lines in the boxes represent the upper (75th) and lower (25th) quartiles and median values, respectively. The whiskers indicate the 95% confidence intervals. The experiments were repeated independently at least three times, with similar results. The n values in the figure indicate the numbers of biologically independent root hair samples.

### Inhibition of PtdIns(3,5)P_2_ production impairs the plasma membrane localization of SYP123, but not that of SYP132, in root hair cells

Next, we tested whether the PM localization of SYP123 and SYP132 in root hair cells would be affected by a reduction in PtdIns(3,5)P_2_ synthesis. In the presence of YM201636, the PM localization of GFP-SYP123 was completely abolished. Instead, GFP-SYP123 fluorescence was observed as punctate structures in the cytosol (Fig. 3A, 3C). The PM localization of GFP-SYP132 was unaffected by treatment with YM201636 (Fig. 3B). Estradiol (1 μM)-induced *FAB1A/B* knockdown caused similar changes in the localization of GFP-SYP123, but not that of GFP-SYP132 (*SI Appendix*, Fig. S2). YM201636-induced cytosolic punctate structures were partially merged with the late-endosomal markers FAB1 and VAMP727 and the trans-Golgi network marker SYP43, indicating that SYP123 localizes to FAB1/VAMP727-positive late endosomes and the trans-Golgi network when FAB1 function is compromised (Fig. 3D). Taken together, these findings indicated that the PM localization of SYP123, but not that of SYP132, is regulated by FAB1 and/or its product PtdIns(3,5)P_2_.

**Fig. 3.**
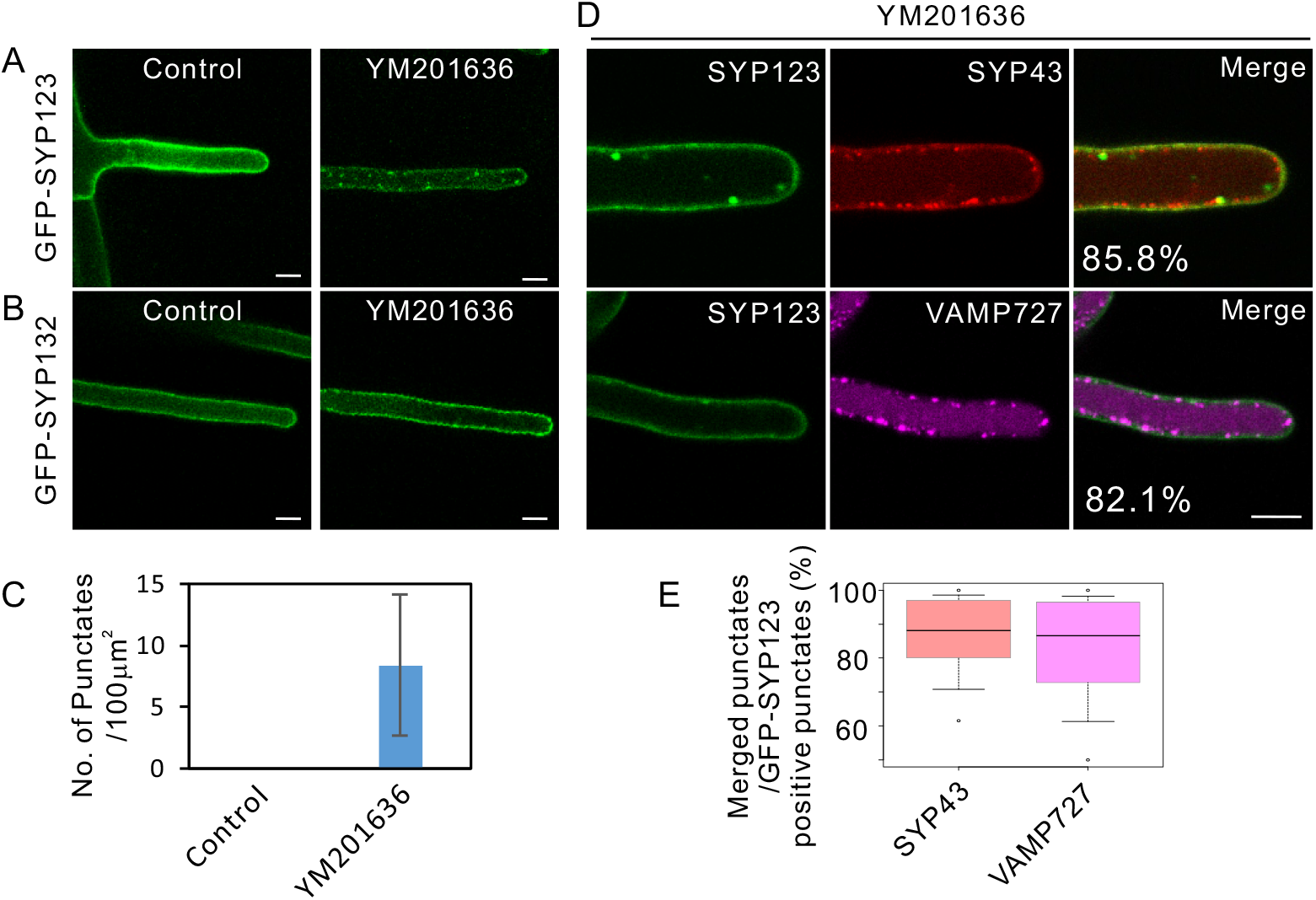
GFP-SYP123, but not GFP-SYP132, relocalizes from the PM to the endosomal compartments upon inhibition of PtdIns(3,5)P_2_ synthesis. Localization of SYP123 (A) and SYP132 (B) in the presence or absence of YM201636. (C) Number of punctate GFP-SYP123 signals in root hair cells in the presence or absence of YM201636. (D) Subcellular localization of GFP-SYP123 in various fluorescent organelle marker lines in the presence of YM201636. Bars = 10 □m. (E) Percentage of merged punctates in (D).

### VAMP727 forms a SNARE complex only with SYP123 on the PM in the root hair shank

The partial colocalization of SYP123 with VAMP727 in the presence of YM201636 implied that they may form a SNARE complex near the PM. To test this possibility, we conducted a bimolecular fluorescence complementation assay to confirm the molecular interaction between SYP123 and VAMP721 or VAMP727 in tobacco epidermal cells. Transient co-expression of the N-terminal fragment of YFP fused with SYP123 (YFPN-SYP123) or SYP132 (YFPN-SYP132) and the C-terminal fragment of YFP fused with VAMP721 (YFPC-VAMP721) or VAMP727 (YFPC-VAMP727) in *N. benthamiana* leaves resulted in fluorescence complementation in combination of YFPN-SYP123 with YFPC-VAMP721 or YFPC-VAMP727, or of YFPN-SYP132 with VAMP721 (Fig. 4A). We further confirmed the interaction between SYP123 and VAMP727 using *in-vivo* co-immunoprecipitation analysis which corroborated that SYP123, but not SYP132, specifically interacts with VAMP727 (Fig. 4B). Taken together, these data suggested that VAMP727 specifically forms a SNARE complex with SYP123 in root hair cells.

**Fig. 4.**
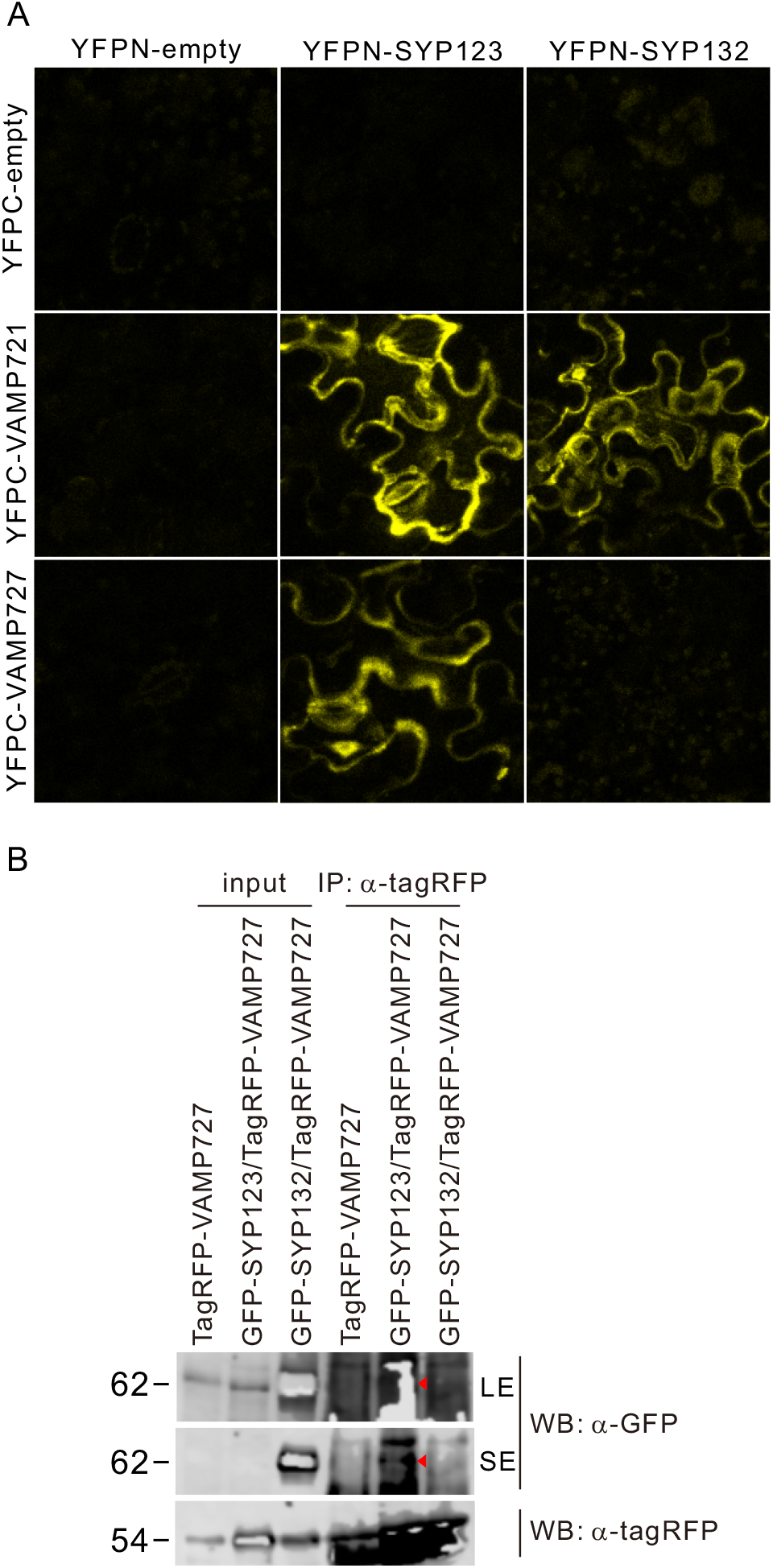
SYP123 physically interacts with VAMP727. (A) Bimolecular fluorescence complementation analysis of SYP123 or SYP132 and VAMP721 or VAMP727 in *N. benthamiana* epidermal cells. Vectors containing the indicated constructs were transiently co-expressed into *N. benthamiana* leaves by Agrobacterium-mediated transformation. Bar = 10 μm. (B) Co-immunoprecipitation assay to examine the specific interaction between SYP123 and VAMP727 in Arabidopsis. Total proteins were extracted from 7-day-old seedlings and immunoprecipitated with anti-TagRFP antibody followed by western blot detection using anti-GFP or anti-TagRFP antibodies. Representative results of three biological replicates with similar results are shown.

To verify the co-localization of SYP123 and VAMP727 on the root hair PM, we observed the PM surface localization of SYP123 and VAMP727 using GFP-SYP123 and Tag-red fluorescent protein (RFP)-VAMP727 co-expression lines. Confocal laser-scanning microscopy revealed that dots of GFP-SYP123 and TagRFP-VAMP727 partially co-localized on the surface of or beneath the PM in the root hair subapical region (Fig. 5A, *SI Appendix*, Movie S3), but not in the cytosol (Fig. 5B). We further confirmed the colocalization of GFP-SYP123 and TagRFP-VAMP727 on the PM by using total internal reflection fluorescence microscopy, which enables observation of the PM surface ^19,29^. Amorphous GFP-SYP123 foci were attached to and moved along with mobile TagRFP-VAMP727 larger dot structures on the PM in the root hair shank. Notably, the globular fluorescence of TagRFP-VAMP727 was surrounded by amorphous GFP-SYP123 fluorescence on the PM surface (Fig. 5C, *SI Appendix*, Movie S4). Collectively, these data suggested that SYP123 on the PM is associated with mobile VAMP727-positive endosomes near or on the PM in the root hair shank.

**Fig. 5.**
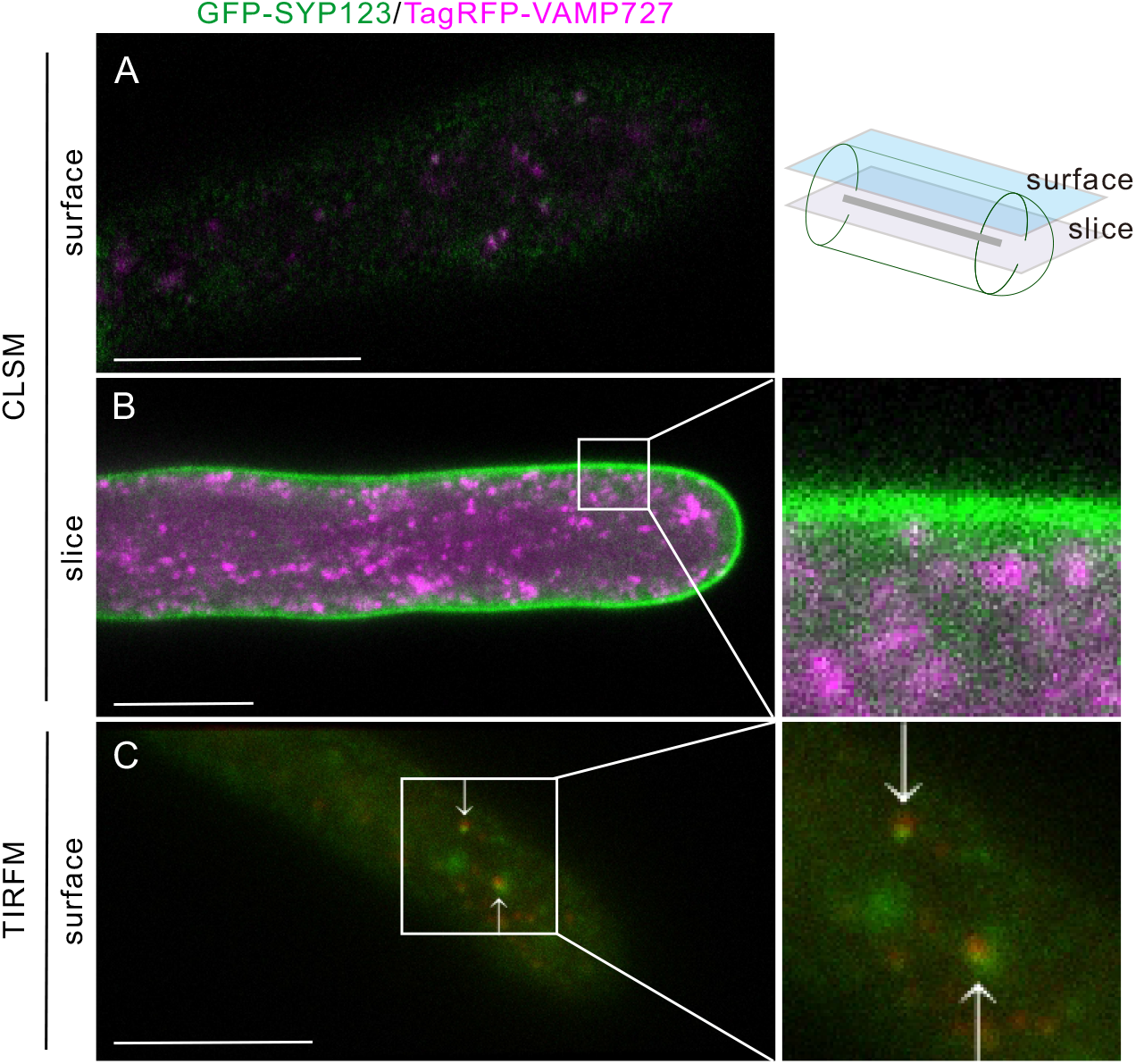
VAMP727-positive endosomes interact with SYP123 on particular domains of the PM in the root hair shank. Co-localization of GFP-SYP123 and TagRFP-VAMP721 on the PM surface (A) or a root hair cell (B). Bars = 10 □m. (C) Total internal reflection fluorescence microscopic image of the PM surface. Fluorescence of GFP-SYP123 (green) and TagRFP-VAMP727 (red) are shown. Bars = 10 □m.

### SYP123 and VAMP727 are required for cell wall stiffness of the root hair shank

We previously reported that reduced *FAB1* expression results in wavy and broader root hairs ^23^. Root hairs of the *syp123* and *vamp727* mutants were broadened and shortened, and slightly bent (Fig. 2A, 6B). Therefore, we hypothesized that loss of function of SYP123 and VAMP727 might impair root hair shank stiffness, as does loss of function of FAB1. To test this hypothesis, we directly measured the stiffness and observed the root hair surface shape in *syp123* and *vamp727* mutants using atomic force microscopy (AFM). Root hair shanks of *syp123* and *vamp727* plants were significantly weaker than those of WT plants (median shank stiffness of WT, *syp123*, and *vamp727* was 7.68 MPa, 3.62 MPa, and 4.27 MPa, respectively, Fig. 6C). Intriguingly, while WT root hairs had a rough surface, those of the *syp123* and *vamp727* mutants had a smoother surface (Fig. 6D). These findings indicated that the *syp123* and *vamp727* mutants have alterations in the stiffness and cell wall surface structure of the root hair shank.

**Fig. 6.**
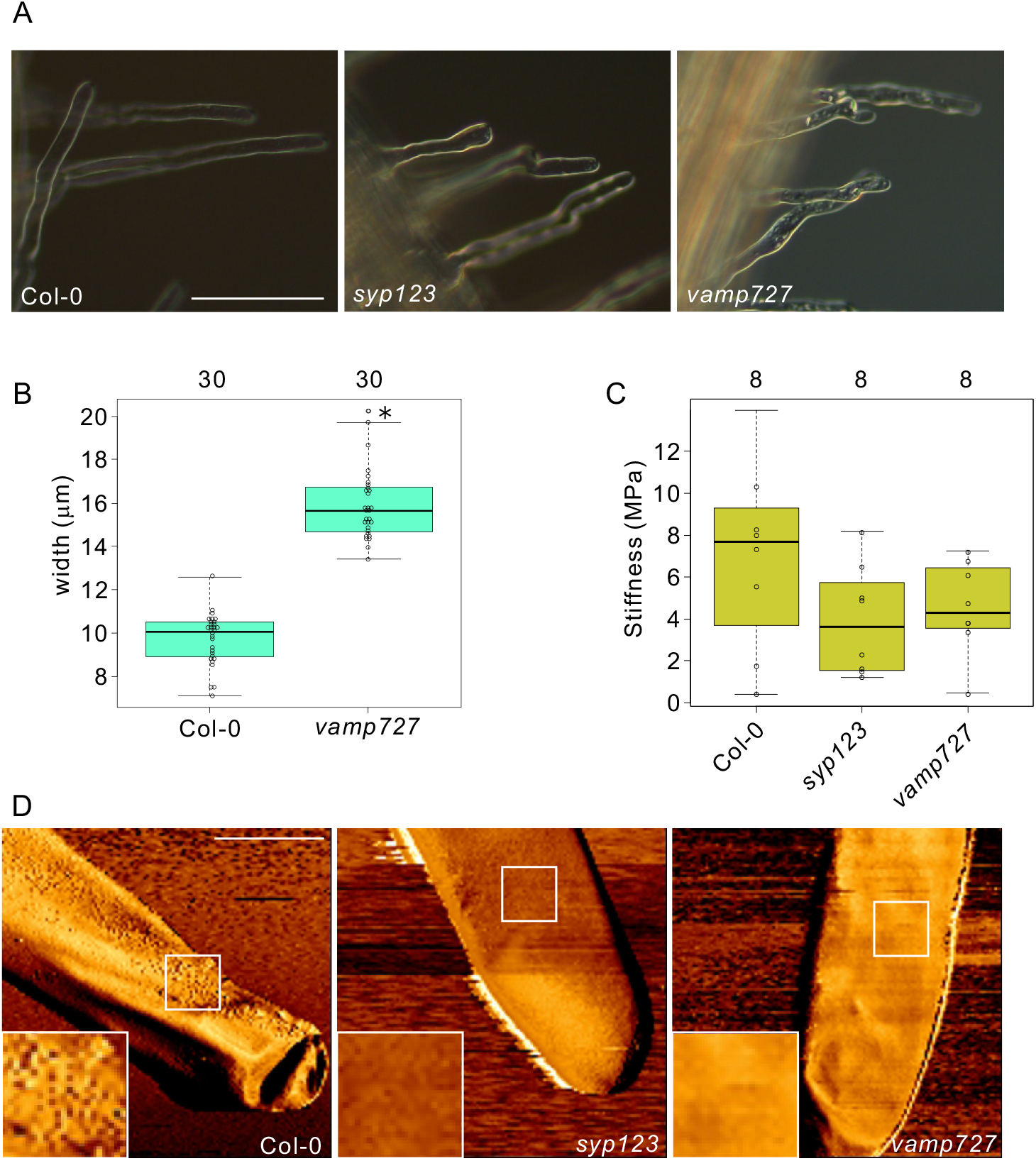
The *syp123* and *vamp727* mutants have weaker root hair shanks. (A) Root hair shapes of WT, *syp123*, and *vamp727*. Bars = 100 mm. (B) Root hair width of WT and *vamp727* mutant plants. Significant differences (\**P* < 0.01) among genotypes were determined for each condition by the Mann-Whitney U-test. (C) Box plots showing root hair shank stiffness. Cell wall stiffness was measured directly by AFM. In the box and-whisker plots, the boxes and solid lines in the boxes show the upper (75th) and lower (25th) quartiles and median values, respectively. The whiskers indicate the 95% confidence intervals. (D) Root hair surface structures as observed by AFM. The squares in the figure are enlarged. Bar = 10 □m.

### SYP123 mediates xylan deposition for inner cell wall construction in the root hair shank

As we previously reported that knockdown of *FAB1* or inhibition of PI(3,5)P_2_ synthesis caused defects in xylan deposition on the cell wall and cortical microtubule construction in the root hair shank ^23^, we next investigated whether SYP123 and VAMP727 are involved in the secretion of inner secondary cell wall components by measuring xylan deposition in the root hair cell wall in *syp123* and *vamp727* mutants by immunostaining with an anti-xylan antibody, LM10 ^30^. The fluorescence intensity of LM10 was significantly weaker in *syp123* and *vamp727* plants than in WT plants (Fig. 7A and 7B). The content of lignin, another cell well component, was also decreased in the root hairs of *syp123* (*SI Appendix*, Fig. S3). In contrast, the fluorescence of an anti-xyloglucan antibody, LM15 ^31^, was unchanged in the mutants (*SI Appendix*, Fig. S4).

**Fig. 7.**
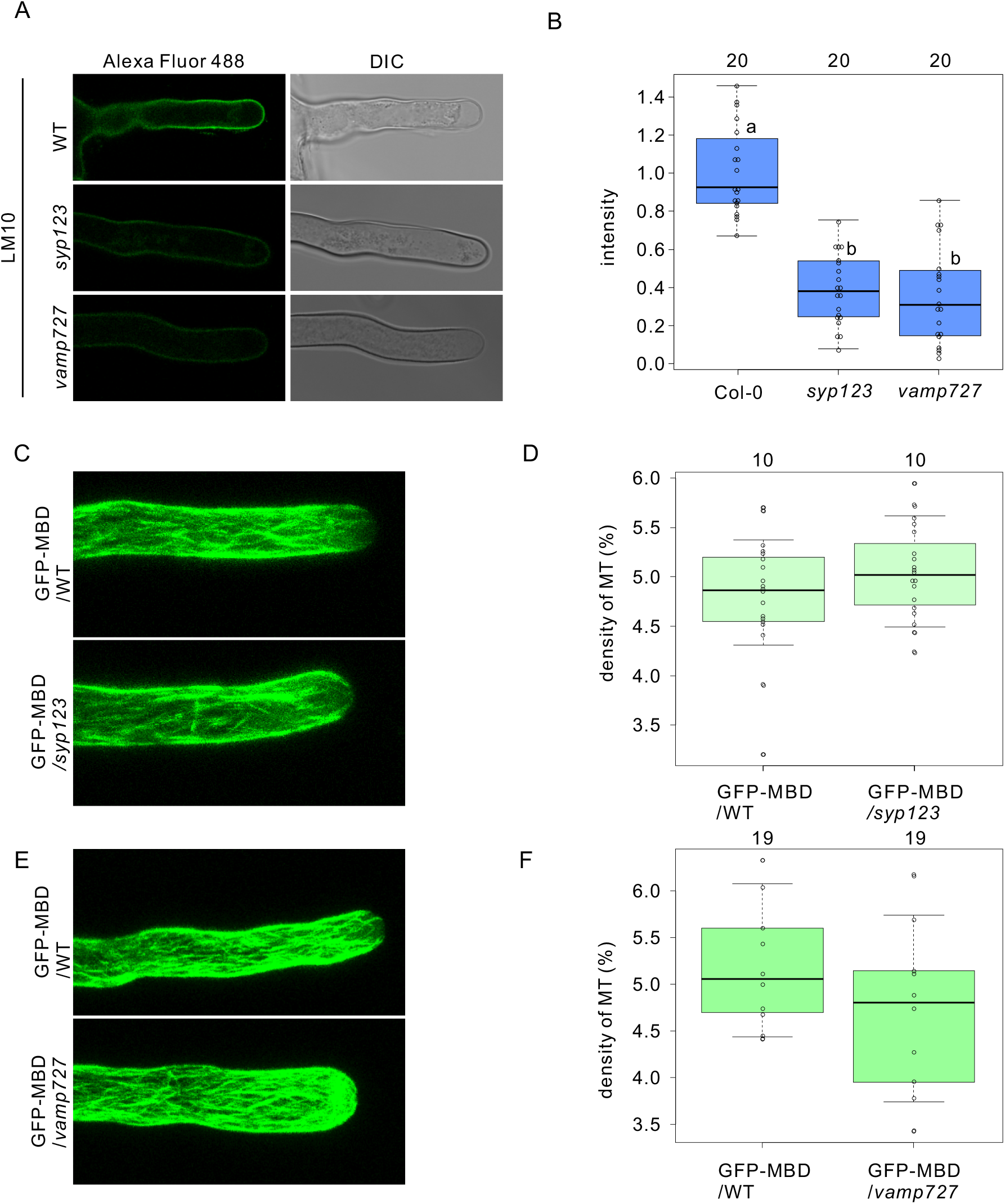
Inner cell walls, but not cortical microtubule structures in the root hair shank, are impaired in the *syp123* and *vamp727* mutants. (A) Indirect immunofluorescence assay using LM10 (anti-xylan rat monoclonal antibody) with root hairs of 5-day-old seedlings of WT, *syp123*, and *vamp727*. (B) Box- and-whisker plots showing the fluorescence intensity in (A) (n = 20 root hairs for each genotype for three biological replicates). The boxes and solid lines in the boxes show the upper (75th) and lower (25th) quartiles and median values, respectively. Groups that have different letters are significantly different from each other (*P* < 0.05, Wilcoxon and Steel–Dwass tests). (C) 3D projection images of GFP-MBD in the growing root hair of 5-day-old seedlings of GFP-MBD in the WT, *syp123*, and *vamp727* backgrounds. (D) Microtubule density in root hairs shown in (C). *P* < 0.01, two-tailed Student’s *t*-test, three independent experiments. n, number of root hairs scored for each genotype or transgenic line.

### Cortical microtubule organization of the root hair is unaltered in *syp123* and *vamp727* mutants

We previously reported that reduced FAB1 activity or PtdIns(3,5)P_2_ production caused defective organization of cortical microtubules, affecting microtubule plus-end stability (Hirano et al., 2018). Therefore, we next examined the cortical microtubule organization of root hair cells using a GFP-microtubule binding domain (MBD) expression line ^32^. Unlike in the case of loss of function of *FAB1* or inhibition of PtdIns(3,5)P_2_ synthesis, the cortical microtubule array of root hair cells appeared normal in the *syp123* (Fig. 7C and 7D) and *vamp727* (Fig. 7E and 7F) mutants, suggesting that SYP123 and VAMP727 are not involved in microtubule organization in root hair cells.

### Computational model of root hair morphology in *syp123* and *vamp727* mutants

While reduced FAB1 function induces wavy root hairs, the *syp123* and *vamp727* mutants revealed no wavy and broadened root hair morphology, despite the lower cell wall stiffness in the root hair shank. This suggests that the wavy root hair morphology is not simply caused by weakening of the cell wall at the root hair shank, but rather by disruption of the cortical microtubule structure in the root hair shank. To evaluate this hypothesis, we adopted a previously reported ^23^ computational model to describe the root hair growth process, with minor modifications. The phenomenological model made the following assumptions based on our observations: (i) root hair growth occurs at the apex, (ii) turgor exerts pressure on the entire root hair cell, (iii) microtubules stabilize root hair shape, (iv) cell wall stiffness of the shank restricts width, and (v) agarose gel culture substrate mechanically represses root hair extension. Tip growth was described by extending the cell wall gradually at the growth point, which is defined as the point at the outermost apex. Cell wall growth was implemented in a two-dimensional framework so that root hair extension depended on root hair width due to the constant increment in surface area of the tip in the three-dimensional structure (see details in Materials and methods). We adapted the experimental data for the *syp123* mutant in this study to fit the assumptions of the computational model. The computational model comprised four parameters, *σ*, *k^tr^*. *k^bend^*, and *k^pip2^*, of which *σ*is the stiffness against mechanical deformation, *k^tr^* is the strength of the action restricting width, *k^bend^* is the bending rigidity, and *k^pip2^* is the relative strength of the effect of PtdIns(3,5)P_2_ on the WT. We considered that *σ* and *k^tr^* represent effects of cell wall construction, and *k^bend^* represents the effect of cortical microtubule formation. The parameter *k^pip2^* controls the three mechanical properties; thus, a *k^pip2^* reduction has a similar effect as a coinciding reduction in *σ*, *k^tr^*, and *k^bend^*.

We assumed that the weakened cell wall structure in the *syp123* mutant reduced the stiffness and width restriction, as shown in Fig. 8C. Accordingly*, k^tr^* = 0.05 and *σ* = 0.5 were set for *syp123*, in contrast to *k^tr^* = 0.1 and *σ* = 1 for the WT. For the *FAB1* knockdown phenotype, it was assumed that the bending rigidity, which is considered to depend on cortical microtubules, was reduced in addition to stiffness and width restriction. These assumptions for *FAB1* downregulation were described as a PtdIns(3,5)P_2_ reduction implemented by assigning *k^pip2^* = 0.5. and *k^pip2^* = 1 for the WT and *syp123* mutant, respectively.

**Fig. 8.**
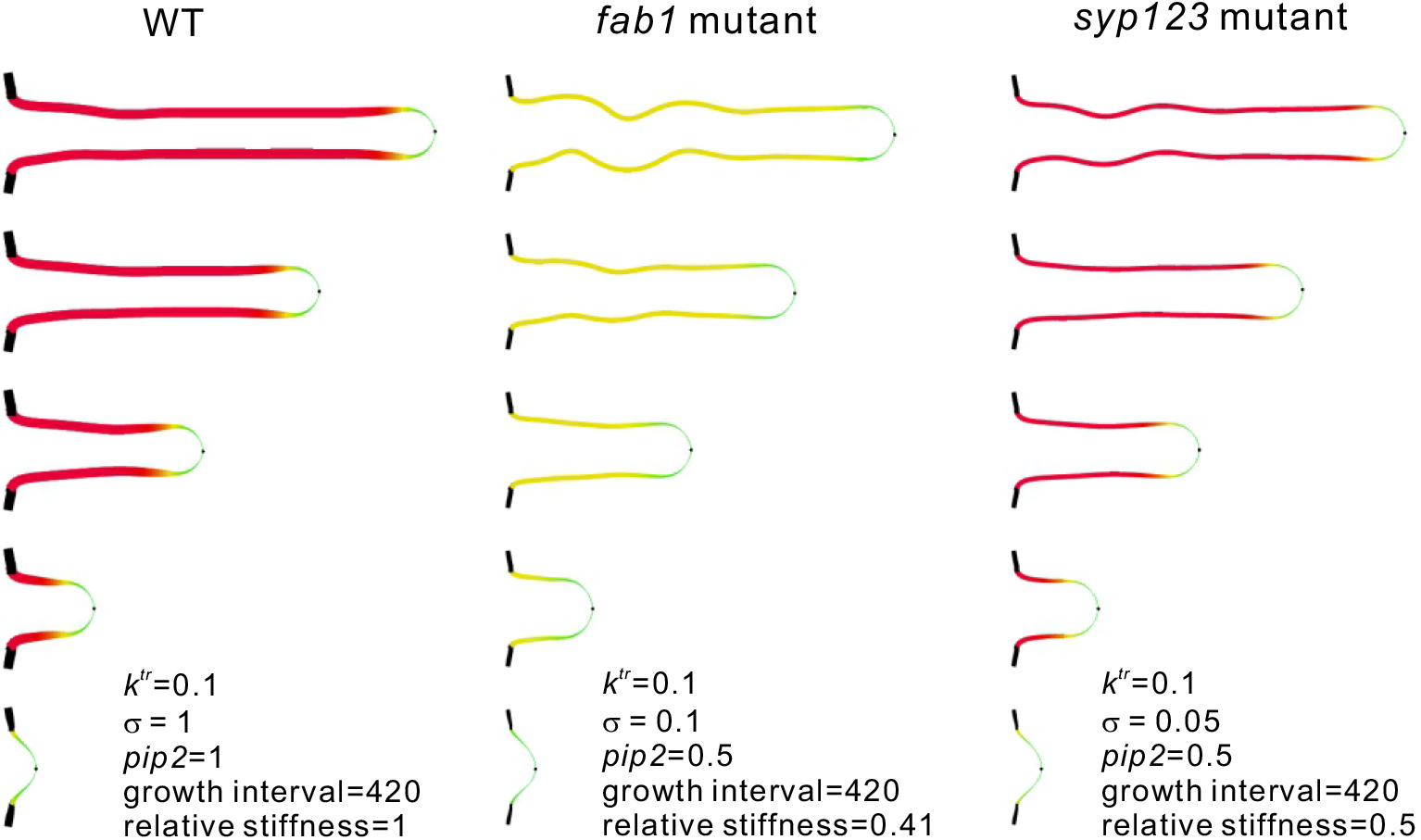
Computational modeling of how mechanical properties regulate root hair morphology. Snap shots of root hair growth for 0, 180,000, 360,000, 540,000, and 720,000 time steps are shown (from bottom to top). Color represents the *pip2*(**x**) values, and line thickness represents the *σ pip2*(**x**) values. (Representation of *k^tr^pip2*(**x**) by line thickness yields similar results.) Black parts on the cell body indicate fixed parts. Black dots at the tip indicate the growth point. Models for WT (left panels), *FAB1* knockdown (KD) (middle panels), and the *syp123* mutant (left panels) are shown.

Figure 8 shows the results of the computational simulation. The root hair morphology depends on cell wall construction (indicated by line thickness) and PtdIns(3,5)P_2_ activity (indicated by color). WT and *FAB1* knockdown model hairs elongated in straight and wavy fashions, respectively (*SI Appendix*, Movie S5, S6). The mechanical properties were reinforced as PtdIns(3,5)P_2_ was activated in the shanks of WT hairs and were kept immature in the *FAB1* knockdown model with low PtdIns(3,5)P_2_ activation. The *FAB1* knockdown model also reproduced the thicker and slightly shorter hairs than those of the WT model, which was in accordance with our observations. The *syp123* mutant model showed slight deformation owing to defects in stiffness and width restriction; however, no wavy morphology was observed (*SI Appendix*, Movie S7). Thus, these results suggested that the wavy phenotype requires a reduction in bending rigidity, which is considered to depend on the cortical microtubule structure.

## Discussion

Re-examination of the localization of GFP-SYP123 in the growing root hair revealed that GFP123 localized in the cytosol rather than in the PM of the root hair tip. PM localization of GFP-SYP123 was observed in the subapical region and the shank of the root hair, but not in the root hair tip, during root hair elongation. As the subapical localization of SYP123 was consistent with that of SYP124 and SYP125 during pollen tube growth ^24–26^, we concluded that SYP123 is mainly localized in the subapical region of the growing root hair, similar to the subapical localization of SYP124 and SYP125 in pollen tubes. This raised the question: what is the precise function of SYP123 in root hair morphogenesis?

We showed that the *syp123* mutant had shortened and broadened root hairs (Fig. 2), but the surface area was unchanged by impairment of SYP123 function, suggesting that *de novo* secretion of root hair PM materials, including primary cell wall components, lipids, and proteins, was not impaired by *syp123* mutation. These results implied that SYP123-dependent secretion is not involved in the delivery of PM and cell wall components necessary for root hair growth. We also found that the inner cell wall components and xylan deposition were severely impaired, thereby lowering the stiffness of the root hair shank. Taken together, these results indicate that SYP123 functions in the secretion of inner cell wall components to harden the root hair shank to maintain the tubular morphology of the root hair.

We demonstrated that VAMP727 also interacts with the root hair-specific SYP12 member, SYP123, in the root hair shank. In addition, we showed that *syp123* and *vamp727* mutants have shortened and broadened root hairs because of weakening of the root hair shank. As xylan deposition was impaired in both *syp123* and *vamp727* mutants, the weakened root hair shank phenotype may be caused by defects in the construction of the inner cell wall structure in the root hair shank. Consistent herewith, deposition of the structural cell wall protein PRP3 and JIM5 antibody-labeled partially methyl-esterified homogalacturonan in the root hair cell wall was impaired in *syp123* mutant and dominant-negative SYP123 plants ^33^. Based on these collective findings, we conclude that the SYP123/VAMP727-mediated late-endosomal trafficking pathway is likely involved in the deposition of cell wall components required for stiffness of the root hair shank, but not for root hair tip growth.

We previously reported that a reduction in FAB1 function results in the formation of wavy root hairs in agarose gel because of disruption of the cortical microtubule structure and weakening of the cell walls of the root hair shank cause mechanical weakness of the shank against the repression force of the gel against the direction of root hair growth ^23^. However, in this study, we hardly observed wavy root hairs in the *syp123* and *vamp727* mutants, although the root hair shank was weakened to the same extent as upon *FAB1* knockdown. These results suggested that the wavy root hair shape may not simply be caused by mechanical weakness of the root hair shank. Therefore, we carried out a computational simulation developed by ^23^, with minor modification. The *syp123* mutant model was slightly deformed owing to defects in stiffness and width, without significant wavy morphology. From the simulation results, we concluded that the wavy phenotype requires a reduction in bending rigidity, which is considered to depend on the cortical microtubule structure, not cell wall rigidity, in the root hair flank. Collectively, our studies imply that two different secretion pathways meditated by SYP123 and SYP132 transport different cell wall components to distinct domains of the PMs of root hair cells. Future studies are needed to elucidate how the different cargo molecules would be transported by these two secretion pathways in plants.

## Methods

### Plant material and growth conditions

*Arabidopsis thaliana* ecotype Columbia-0 (Col) plants and T-DNA insertion mutants in the Col-0 background were used. The *syp123* (CS488587) and *vamp727* (GABI_060G05) mutants and a FAB1A/B artificial microRNA line generated previously ^27^ were used in this study. The transgenic marker lines GFP-SYP123 ^12^, FAB1A-TagRFP ^23^, TagRFP-SYP43 (16), TagRFP-VAMP727 ^16^, SNX1-TagRFP (Jaillais et al., 2006), and 35S::GFP-MBD ^32^ were used in this study. Plants were grown under continuous white light at 22 ± 2°C on 1/2 Murashige and Skoog (MS) medium (1% sucrose, 1.2% agarose). To inhibit FAB1A and FAB1B, plants were grown on 1/2 MS medium supplemented with 1.0 μM YM201636 (Cayman Chemical) for 2 days.

### Measurement of root hair length and microtubule density in root hairs

Root hair length was measured in 5-day-old seedlings, as described by ^34^, with some modifications. Root hairs were digitally photographed using a stereomicroscope at a magnification of 40–50×. The length of five consecutive hairs protruding perpendicularly from each side of the root, for a total of 10 hairs from both sides of the root, was determined using ImageJ. The microtubule density in WT, *syp123*, and *vamp727* root hairs was measured as described previously ^23^.

### Confocal microscopy

Root hairs were imaged using a Leica TCS SP8 laser scanning confocal microscope. For the imaging of GFP/RFP combinations, samples were excited using 488- and 552-nm laser lines in multitrack mode. Signals were collected using a HyD detector (Leica) with emission wavelengths of 500–530 nm for GFP and of 590– 650 nm for TagRFP. Fluorescence intensities were measured using Image Studio software (LI-COR).

### Total internal reflection fluorescence microscopy

Arabidopsis seedlings expressing GFP-SYP123 and TagRFP-VAMP727 were grown under continuous white light at 22 ± 2°C on 1/2 MS medium (1% sucrose, 1.2% agarose) in a 35-mm glass-base dish (Iwaki) for total internal reflection fluorescence microscopy. Growing root hairs were observed under an IX-71 (Olympus) equipped with a UAPON 100x O TIRF lens (Olympus), as described in Fujimoto et al. (2020), with some modifications. GFP and TagRFP were simultaneously excited using 473- and 561-nm lasers, respectively. An FF01-523/610-625 filter (Semrock) was used to remove autofluorescence. Fluorescence emission spectra were separated using a FF560-FDi01-25×36 LP dichroic mirror (Semrock) and filtered through a FF01-523/35 filter (Semrock) for GFP and an ET620/60M filter (Chroma) for TagRFP, using DV2 (Photometrics). Images were acquired using an iXon X3 EMCCD camera (Andor Technology) operated with Metamorph software (Molecular Devices). Each frame was exposed for 150 ms. The images were analyzed using ImageJ.

### Imaging and indentation measurement by AFM

AFM imaging and force curve measurements were conducted using a Nanowizard II (JPK BIOAFM instruments/Bruker) equipped with an inverted optical microscope (Axio Observer D1; Zeiss). A root with root hairs was immobilized on a plastic Petri dish with scotch tape. First, the root hairs were observed by phase-contrast imaging, followed by AFM imaging and force curve measurement. AFM imaging was conducted in tapping mode using an OMCL-AC240TN-R3 cantilever with a resonance frequency of 70 kHz in air, 2 N/m spring constant, and 7 nm tip radius (Olympus). Subsequently, force curves were measured on indicated points on the root hairs using the same cantilever. The force curves were processed and analyzed using the manufacturer’s software. The indentation zones of the approach traces were fitted with Hertz’s model to estimate Young’s moduli.

### Detection of cell wall components

Arabidopsis seedlings were grown vertically on MS medium for 5 days, gently removed from the agarose surface, placed in a Petri dish, and incubated in PBS buffer containing 1.0% (w/v) BSA for 30 min for blocking. After blocking, the seedlings were washed with PBS for 30 min three times and then placed in primary antibody solution that consisted of a 1/10,000 (v/v) dilution of wall polymer-specific antibody (LM10, LM15) in PBS. Plates containing seedlings were gently shaken on a rotator in the dark at room temperature (RT) for 90 min. The seedlings were with PBS three times, blocked (as described above), and washed again. The seedlings were incubated and gently shaken in the dark at RT in 500 μL of a 1/10,000 (v/v) dilution of secondary antibody (anti-rat-Alexa Fluor 488; Thermo Fisher Scientific) in PBS for 90 min. Then, they were washed with PBS for 30 min three times. Roots hairs of 5-day-old Arabidopsis seedlings were stained with basic fuchsin as previously described (Ursache et al., 2018).

### Bimolecular fluorescence complementation assays

The coding regions of *SYP123, VAMP721*, and *VAMP727* were cloned into the Gateway entry vector pENTR-D/TOPO (Invitrogen) and subsequently transferred to the Gateway-compatible binary vectors pBiFP2 and pBiFP3 ^35^. Primers are listed in Supplementary Table S1. *Nicotiana benthamiana* plants were transiently transformed with *Agrobacterium tumefaciens* GV3101 carrying the NYFP-SYP123 and NYFP-SYP132 constructs in combination with CYFP-VAMP721 and CYFP-VAMP727.

### Coimmunoprecipitation and western blotting

Five-day-old seedlings of GFP-SYP123 and a TagRFP-VAMP727 co-expression line were frozen in liquid nitrogen and ground into powder. Proteins were extracted in lysis buffer (50 mM Tris-Cl [pH 7.4], 150 mM NaCl, 1 mM EDTA, 1 mM CaCl2,1% Triton X-100). For immunoprecipitation, 50 □L of Protein G Sepharose (GE Healthcare) was used in accordance with the manufacturer’s instructions. In brief, an anti-TagRFP antibody (Evrogen) was added to the beads. The beads with the antibody were rotated top to bottom overnight at 4°C. The next day, the beads were washed gently and suspended in sodium dodecyl sulfate loading dye. The sample was heated at 95°C for 5 min, and the isolated proteins were analyzed by western blotting using anti-GFP primary antibody (1:2,500, Novus Biologicals) and goat anti-rabbit IgG-horseradish peroxidase secondary antibody (1:5,000, GE Healthcare).

### Statistics

In box-and-whisker plots, the boxes and solid lines in the boxes indicate the upper (75th) and lower (25th) quartiles and median values, respectively. The whiskers indicate the 95% confidence intervals. Unless stated otherwise, the significance of differences between two samples was evaluated by a Mann–Whitney U-test. In the case of multiple comparisons, significant differences (*P* < 0.01; indicated by different letters) among genotypes were determined for each condition by Wilcoxon and Steel–Dwass tests.

### Computational model

The mechanical model describing the root hair growth process was modified based on our previous study ^23^. The basic concept of the model is to draw the shape of a root hair by a sequence of particles spaced evenly in a two-dimensional space (Takigawa-Imamura 2015). Each particle is assumed to move depending on interaction forces as follows:

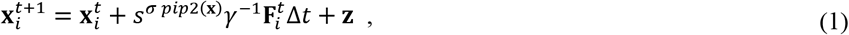

where 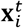 is the position of the *i*-th particle, 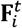 is the sum of the forces working on the particle, **z** is the random vector representing the noisy fluctuation of the cellular behavior, *t* is the time step, and Δ*t* is the time-step size. The parameter *γ* corresponds to the basic stiffness of the cell wall against mechanical deformation. The parameters *s* and *σ* determine the enhanced stiffness by cell wall construction in the shank (0 < s < 1, 0 ≤ *σ* ≤ 1). We set *γ* = 8, *s* = 0.25, and *σ* = 1. The value *pip2*(**x**) represents the local PI(3,5)P_2_ concentration, which ranges from 0 (at the tip) to *k^pip2^* (in the shank region) ^23^. We set *k^pip2^* = 1. The reduction in FAB1 expression was described by *k^pip2^* = 0.5.

The sum of the forces 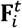 calculated at each time step was assigned as follows:

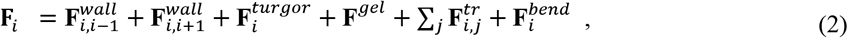

where 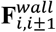 is the interaction between adjacent particles for sustaining a constant space, 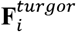 is the turgor pressure pushing each particle outward, and **F**^*gel*^ is the force exerted by the agarose gel that represses root hair extension ^23^. 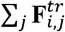 is the summation of forces restricting root hair width, where *j* is chosen so that the vector between the *i*-th and *j*-th particles is nearly vertical to the cell wall (see the definition of 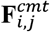 in Hirano et al. (2018). We considered that cell wall construction in the shank restricts root hair width. Cell wall construction was assumed to depend on PI(3,5)P_2_, and 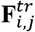 was defined as follows:

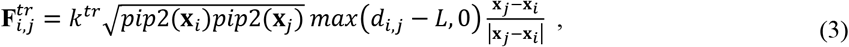

where *k^tr^* is the basic strength of the restriction force, *max*(*a,b*) is the function that takes the larger between *a* and *b*, and *L* = 4.3 is the prescribed value to adjust the restriction force.

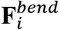 denotes the bending rigidity, which was assumed to be caused by cortical microtubule formation. Cortical microtubule formation was assumed to depend on PI(3,5)P_2_, and 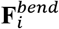 was defined as follows:

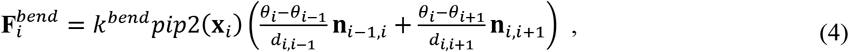

where *k^bend^* is the basic strength of bending rigidity. The variables and vector *Θ_i_, d_i,j_*, and **n**_*i,j*_ describe the shape around the *i*-th particle (see the definitions in Hirano et al., 2018). We set *k^bend^* = 4.

The numerical calculation was initiated where a small bud sprouting from the epidermal cell body was depicted (Fig. 8, lowest panels). Root hair growth was described by inserting a new particle at the outermost tip ^23^. To control the insertion interval, the growth rate *r* = 4×(tip radius) × (particle diameter) ÷ (insertion interval), was defined by considering the constant increment in surface area per time step in a threedimensional structure. The insertion of new particles occurs depending on *r* = 0.0019, particle diameter 0.4, and width near the root tip measured each time step during the numerical calculation. The number of particles in the initial state was 42, which increased to 284 (WT), 250 (FAB1 KD), and 272 (*syp123*) after 720,000 time steps.

## Acknowledgments

We thank Ms. Mina Yamamoto for measuring of the root hair length of WT and *syp123* mutant plants. This work was supported by JSPS KAKENHI J19H00933 and JP16H06280 to M.H.S., 20K05962 and a Grant for Basic Science Research Projects from The Sumitomo Foundation to T. Hirano.

## Data Availability

All study data are included in the article and *SI Appendix*.

## Author Contributions

T. Hirano and M. H. S. conceived and designed the study. T. Hirano, M. Y., K. E., T. N., H. K., T. U., and T. Higaki performed the experiments and analyzed the data. H.T.-I performed the mathematical modeling. M.H.S. designed the experiments and supervised the work; M.H.S., T. Hirano, and H. T.-I. wrote the paper.

## SI Appendix

### Legends for supplemental figures

**Fig. S1.**
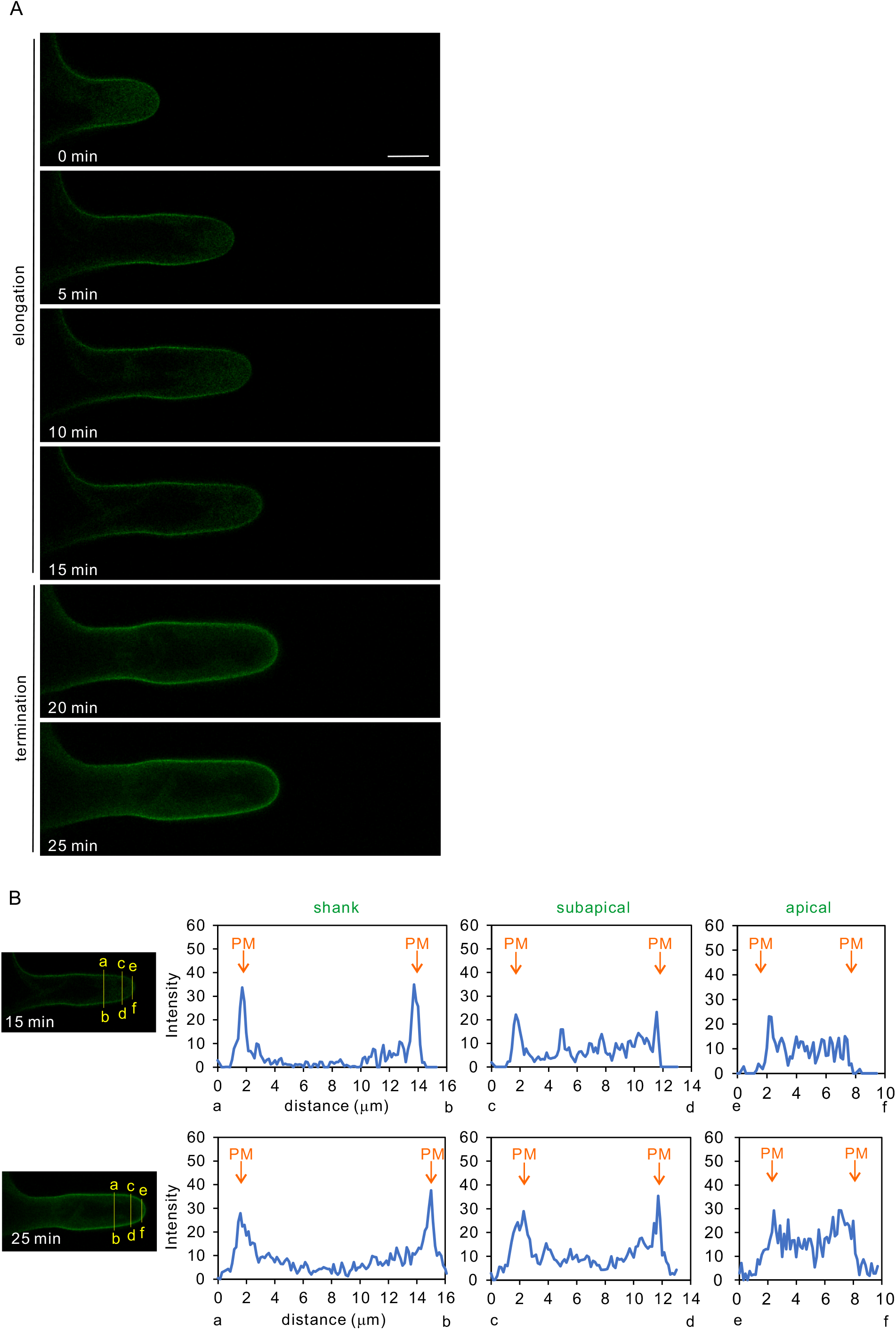
Changes in the localization of GFP-SYP123 on the plasma membrane (PM) in the root hair during elongation and termination stages. (A) Time-lapse images of GFP-SYP123 localization in the elongation and termination stages of a root hair was observed by confocal laser scanning microscopy in 5-min intervals. Bar = 10 μm. (B) GFP-SYP123 intensity profiles in the root hair shank [(a)–(b)], subapical [(c)–(d)], and apical [(e)–(f)] regions at elapsed times of 15 or 25 min, as indicated in (A), are shown.

**Fig. S2.**
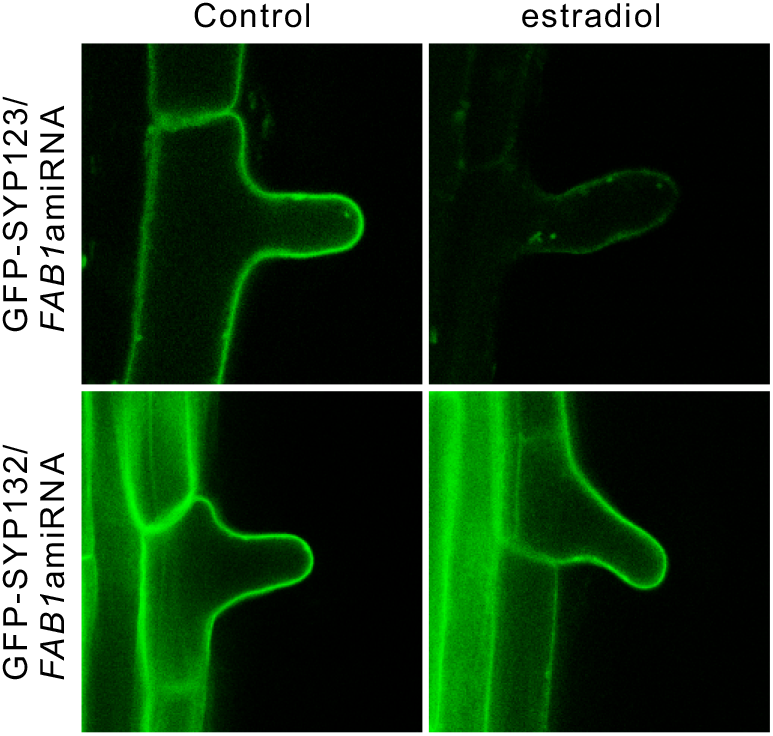
PM localization of SYP123, but not SYP132, depends on the FAB1 late endosome-mediated pathway. GFP-SYP123 (upper columns) or GFP-SYP132 (lower columns) in the background of a FAB1 artificial microRNA line, in the absence (right columns) or presence (left columns) of 1 □M estradiol. Bars = 1 □m.

**Fig. S3.**
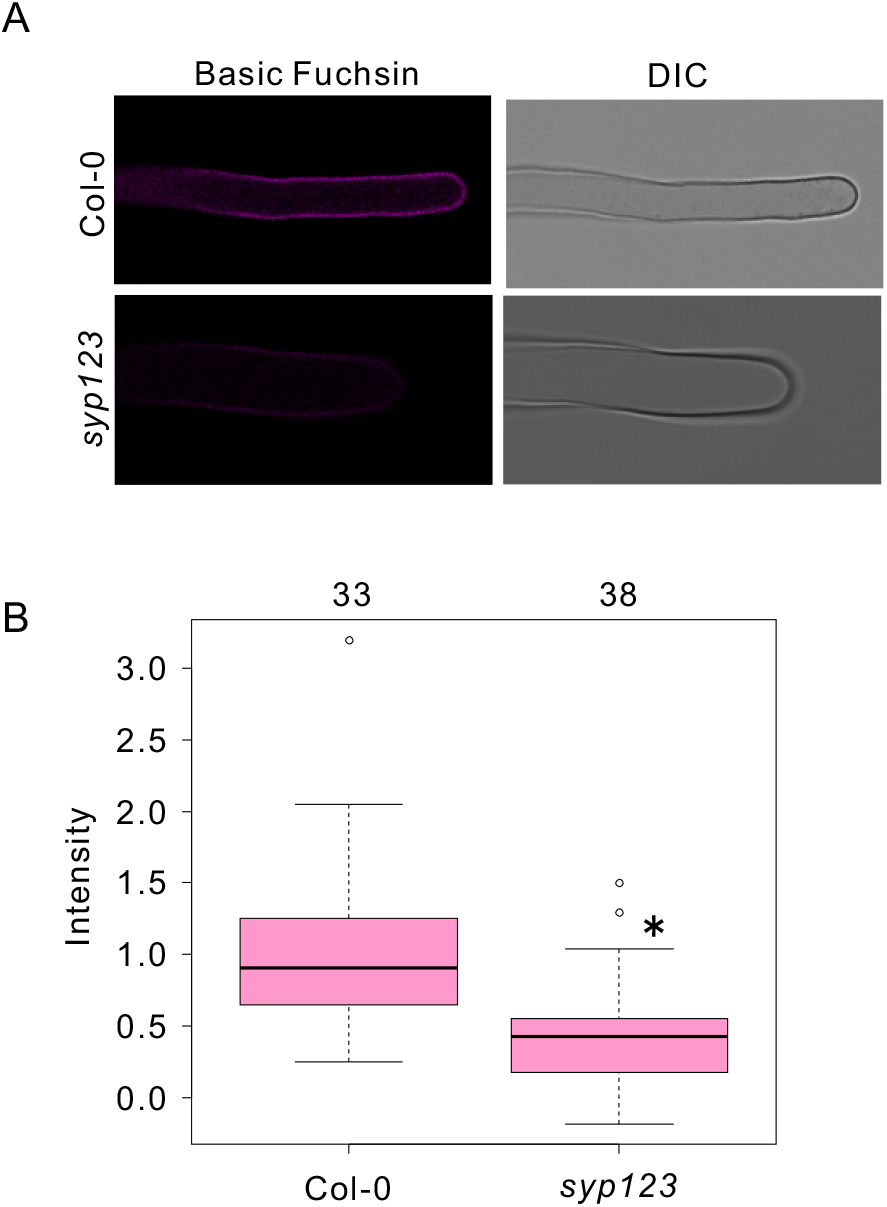
Lignin deposition is impaired in the root hair shank of *syp123*. Images of wild-type (WT) and *syp123* mutant root hairs stained with basic fuchsin (A), and fluorescence intensity profiles of basic fuchsin staining (B). \**P* < 0.05 vs. WT, Mann-Whitney U test. In the box and-whisker plots, the boxes and solid lines in the boxes show the upper (75th) and lower (25th) quartiles and median values, respectively. The whiskers indicate the 95% confidence intervals. The experiments were repeated independently at least three times, with similar results. The n values in the figure indicate the numbers of biologically independent root hair samples assessed.

**Fig. S4.**
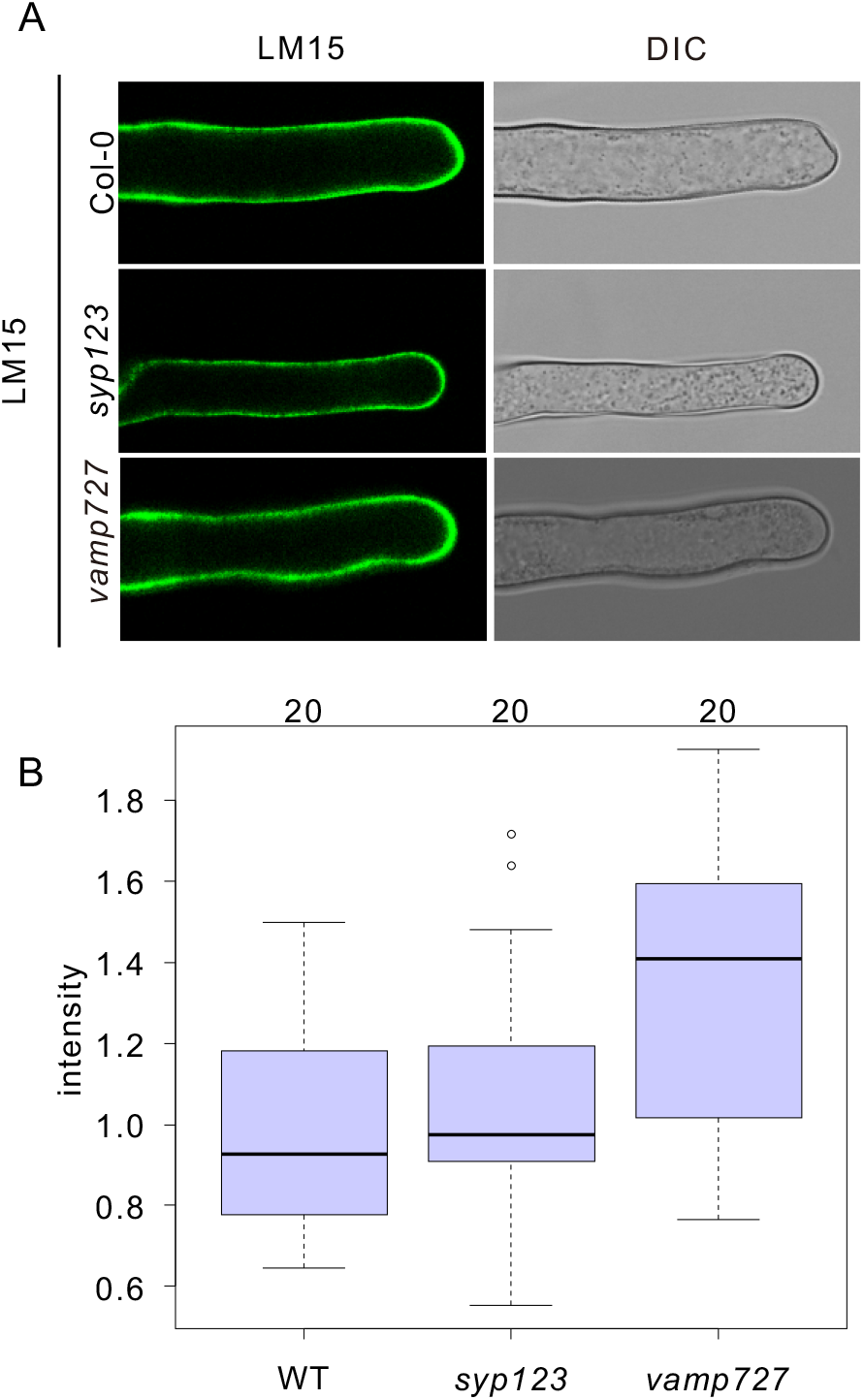
(A) Cell wall of a root hair stained with the anti-xyloglucan antibody, LM15, and Alexa Fluor 488-conjugated anti-rat secondary antibody in WT, *syp123*, and *vamp727* plants. (B) Box and-whisker plots showing fluorescence intensity values. n = 20 root hairs for each genotype. Three biological replicates were included. Boxes and solid lines in the boxes show the upper (75th) and lower (25th) quartiles and median values, respectively. Mutants were compared with Col-0 using the Wilcoxon and Steel–Dwass tests. *P* < 0.05.

**Fig. S5.**
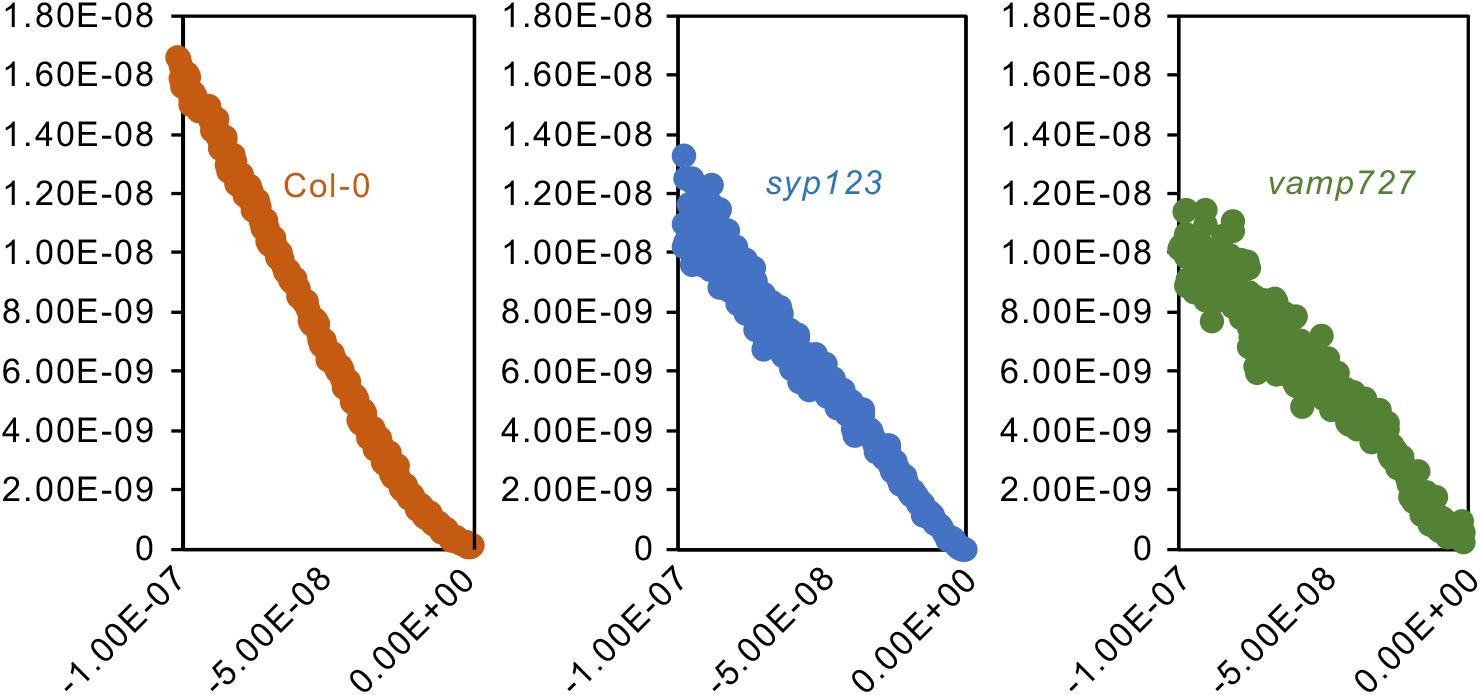
Force curves of atomic force microscopic measurement of Col-0, *syp123*, and *vamp727* root hairs.

### Legends for Supplemental movies

Movie S1. Localization dynamics of GFP-SYP123 in elongating root hair. Time interval is 5 min.

Movie S2. Changes in localization dynamics of GFP-SYP123 during termination of root hair elongation. Time interval is 5 min.

Movie S3. Surface localization of GFP-SYP123 and TagRFP-VAMP727 on root hair shank. The images of confocal scanning laser microscopy are shown. Time interval is 850 msec.

Movie S4. TIRFM images of GFP-SYP123 and TagRFP-VAMP727 on the PM of root hair shank. Time interval is 150 ms.

Movie S4. Computer simulation of the root hair of WT in Fig. 8.

Movie S5. Computer simulation of the root hair of *fab1* knockdown mutant in Fig. 8.

Movie S6. Computer simulation of the root hair of *syp123* mutant in Fig. 8.

